# Evidence for the Progressive Improvement of All-Atom Force Fields in Reproducing Local Conformational Preferences of Flexible Peptides

**DOI:** 10.64898/2025.12.22.695171

**Authors:** Javier González-Delgado, Juan Cortés, Alessandro Barducci, Matteo Paloni

## Abstract

Classical protein force fields are widely used to probe the conformational properties of intrinsically disordered regions, yet their accuracy in reproducing local structural preferences remains uneven. We evaluated seven Amber and CHARMM force fields across three generations using molecular dynamics simulations of glycine–X–glycine tripeptides, with guest residues that span diverse physicochemical properties. Conformational ensembles were compared against distributions of conformations extracted from the crystallographic structures in the Protein Data Bank, and a statistical model derived from NMR observables. Analysis of secondary structure populations and Ramachandran distributions analyzed via Wasserstein distances reveals a clear historical progression. Early models display strong helical bias, intermediate ones approach Protein Data Bank trends, and recent versions shift toward solution-like ensembles dominated by polyproline II structure. None of the force fields fully captures the experimental distributions, although recent models show marked improvement over earlier generations. The remaining discrepancies point to specific aspects of local structure that still require tuning, while the overall progress underscores a steady trajectory toward more reliable descriptions of disordered peptides.

---

The accuracy of the parameters used in molecular dynamics (MD) simulations, *i.e.* the force field (FF), determines their ability to obtain atomistic insight into the conformational dynamics of biomolecular systems and to capture both global and local structural tendencies. This is essential for characterizing intrinsically disordered regions (IDRs), which lack stable secondary structure and require high quality FFs in light of their roles in cellular pathways and disease, for instance as recognition modules^1^, and their rapid exchange between conformations^2^. Considerable work has therefore focused on their structural characterization over the past two decades using both experiments and simulations^3–6^. Atomistic simulations have been used in particular to probe conformational ensembles of IDRs in dilute and dense protein environments^7–9^.

Short peptides provide an ideal setting to study amino acid specific structural properties and to test the reliability of protein FFs by neglecting contributions related to long range order^10^. Different amino acids display clear preferences for *α* helical, *β* strand, and polyproline II (*ppII*) motifs^11–13^. Extensive work has gone into correcting early FF limitations by improving backbone dihedral parameters ^14,15^, rescaling protein solvent interactions^16^, and introducing new water models^17–19^, which affect both local and global structure. Earlier studies used short peptides to identify conformational biases^20^ and evaluate agreement with NMR J-couplings^21,22^. Previous works examined glycine-X-glycine (GXG) tripeptides to test four classical FFs against Ramachandran distributions and J-couplings using a Gaussian mixture model (GMM) representing NMR data^23^. They found that Amber ff19SB reproduced residue specific *ppII* content best among the tested models^23^.

Here we carried out simulations of GXG tripeptides including seven guest amino acids that span size, charge, and hydropathy (Table S1) to compare the performance of protein FFs in reproducing local secondary structures and (*ϕ*,*ψ*) distributions. We evaluated seven Amber and CHARMM force fields developed across the past three decades (Table S2). Conformational propensities from simulations were compared with experimental references, namely (*i*) coil regions extracted from protein structures in the Protein Data Bank (PDB), which have been used to generate accurate ensembles of disordered proteins^24^, (*ii*) a subset of these structures filtered for solvent exposed tripeptides (PDB-Exposed), and (*iii*) a GMM derived from NMR data^25^. Details on these references are given in Supplementary Methods. We first examined the secondary structure propensities of the central residue in the three experimental and all simulated data sets by measuring the populations of *α*, *β*, and *ppII* regions^20^ (Fig. 1a). These regions cover most of the sampled conformations, exceeding ninety percent in nearly all cases. Occupancies for each tripeptide appear in Fig. S1 and in Tables S6 and S7, for experimental and simulated datasets, respectively.

**Figure 1:**
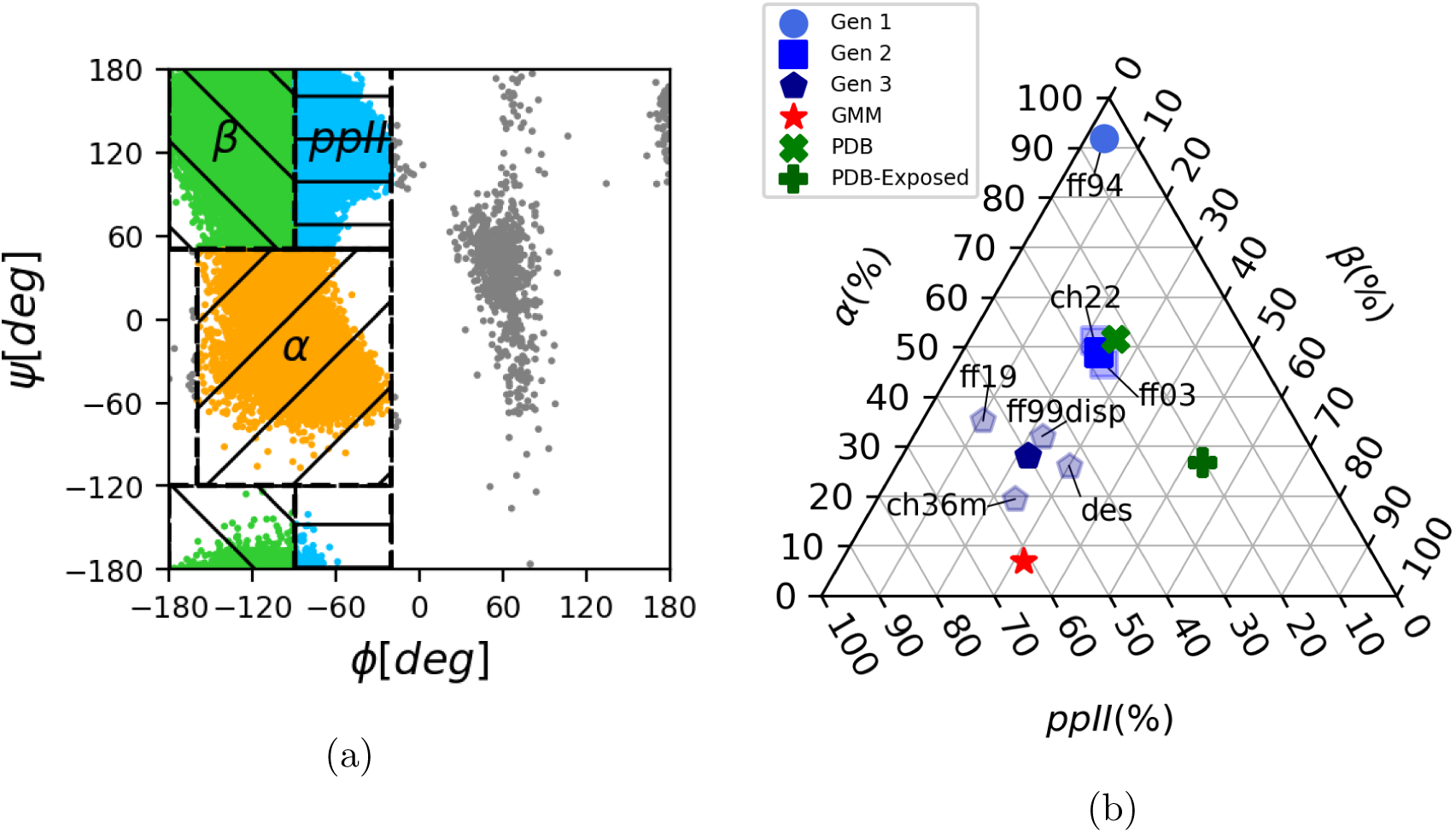
(a) Definition of the secondary structure regions according to Best *et al.*^20^ with a representative assignment of points from MD simulations (GFG tripeptide with DES-Amber). Colors indicate the assignment to the secondary structure regions, gray points indicate conformations that remain unassigned. (b) Triangular plot depicting the secondary structure propensities for each tripeptide, force field and experimental data set. Opaque blue symbols indicate average values for each generation of force fields, transparent blue symbols indicate values for each force field and the marker corresponds to that of the corresponding generation. Symbols in green and red indicate values from the experimental data sets. Gen 1 is composed only by ff94, thus their points overlap. Abbreviations for the force fields are given in Supp. Table 2.

The PDB set shows about 50% *α*, 25% *β*, and 25% *ppII* structure. Residue specific analysis indicates that small residues such as glycine, serine, leucine, and alanine prefer *α* regions, while charged and aromatic residues such as lysine, glutamate, and phenylalanine favor *β* structures. In the PDB-Exposed subset, *ppII* content is almost unchanged, but the *α* population falls to roughly 50% and alanine shows unexpectedly high *β* preference, similar to lysine and phenylalanine. The GMM instead yields dominant *ppII* structure near 60% and about 30% *β* structure with minimal *α* content below 10%, and glycine and alanine reach close to 90% *ppII*.

FFs display substantial variation and naturally group into three chronological classes (Fig. 1b). The earliest model, Amber ff94, which constitutes the *first generation*, strongly favors *α* structure above 90%, consistent with its known bias^20^. The *second generation* of FFs, CHARMM22 and Amber ff03, produces a more balanced pattern close to the PDB values. Later FFs, constituting the *third generation*, aimed to reproduce protein conformations including disordered regions, using 4-point water models. This generation includes CHARMM36m, Amber ff19SB, ff99SB-disp, and DESAmber, and reproduces ensembles closer to solution data from the GMM, with *ppII* around 50%, moderate *α* content near 30%, and *β* near 20-30%. Further analysis is provided in Supplementary Results and reflects steady improvement in parameterization for IDRs.

To quantify agreement between simulated and experimental distributions, we computed the Wasserstein distance (WD) between every pair of (*ϕ*,*ψ*) distributions. This metric is well suited for angular data because it accounts for periodicity, unlike classical metrics, such as total variation, Hellinger or *ℓ_p_* distances, and recent work supports its use for comparing conformational ensembles of flexible proteins at the residue level. ^26,27^ Unlike comparisons based on secondary structure categories, which may discard conformations outside the main populated clusters (for instance the gray points in Fig. 1a), it uses the full distribution and captures differences in peak locations and shapes. Its formal definition is provided in Supplementary Methods.

Figure 2a shows a two dimensional scaling projection based on WDs averaged over replicas and tripeptides. The three experimental references are clearly separated, with the PDB-Exposed subset lying between the other two yet closer to the PDB ensemble. Apart from Amber ff94, all simulation data sets follow a path that spans from the PDB reference toward the GMM, reflecting the historical evolution of FFs and supporting their classification into three generations (Fig. 2b). Amber ff94 falls far from all references, in line with its poor reproduction of local structure, while second generation models overlap with the PDB and third generation models extend toward the GMM. Within this group, Amber variants cluster near the PDB-Exposed subset and CHARMM36m approaches the GMM.

**Figure 2:**
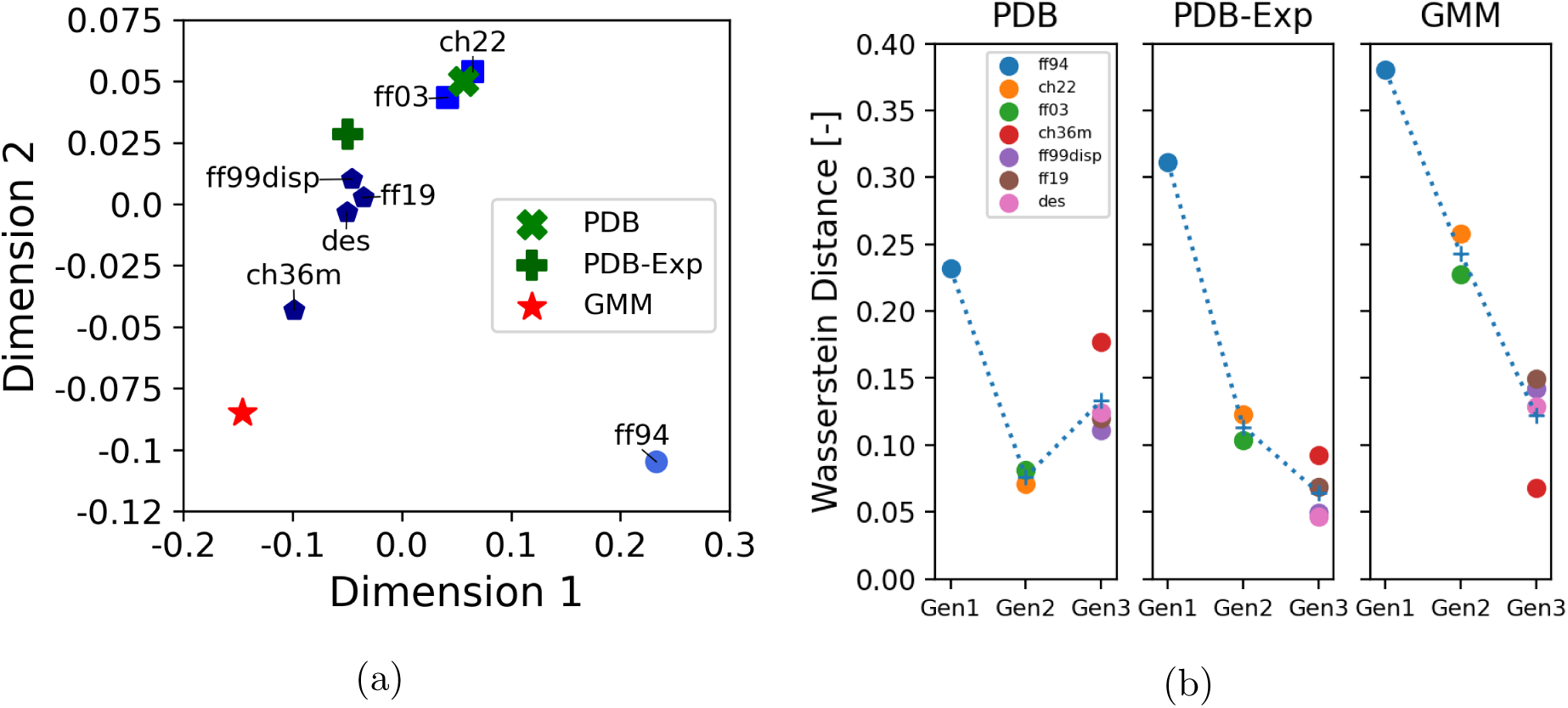
Wasserstein-based multidimensional scaling of (ϕ, ψ) distributions. (a) WDs are averaged over all tripeptides and replicas before performing the scaling. (b) WD between each MD simulation and each experimental data set. Crosses indicate average values for each generation and dotted lines serve as visual guides to indicate the change of the averages with the generations.

This projection refines the picture emerging from the secondary structure analysis. None of the force fields fully reproduces any reference set and deviations from one distribution correspond to improved agreement with another. This behavior indicates that each reference set is affected by biases that limit their accuracy, making it difficult to assess which reference model more faithfully captures tripeptide conformational ensembles in solution. On the one hand, Protein Data Bank derived datasets may be still affected by the molecular environment in the crystallographic structure. On the other hand, GMMs, by construction, are a simplified representation of the complex Ramachandran distributions.

This study provides a detailed evaluation of how classical force fields reproduce local conformational preferences in GXG tripeptides. The combined analysis of secondary structure populations and Wasserstein distances shows clear progress from early biased models to more recent versions that better match experimental trends, though none fully aligns with the reference ensembles. The consistent separation between PDB, PDB-Exposed and GMM data, and the placement of force fields along the path connecting them, highlights the remaining imbalance between conformations typical of folded proteins and those dominant in solution. The clustering of Amber variants near PDB-Exposed and the proximity of CHARMM36m to GMM illustrate this divide.

The approach developed here can be extended in several directions. On the one hand, enhanced sampling methods could allow a systematic exploration of the full peptide space, making it feasible to sample sequences with slow dynamics. On the other hand, the rapid progress in machine learned interaction potentials for biomolecules ^28,29^ promises to offer reference quality distributions for isolated peptides that surpass the present data sets.

## Acknowledgements

This work was supported by the French National Research Agency (ANR) under grant ANR-22-CE45-0003 (CORNFLEX project). This work used the HPC resources of the CALMIP supercomputing center under allocation 2016-P16032.

## Supplementary Information Supplementary Results

### Secondary Structure Populations of Guest Residues from MD Simulations

Different amino acids display characteristic deviations within each force field family. In Amber ff94, nearly all residues populate *α* regions almost exclusively (*>*90%), with minimal variation; only glycine shows a slightly reduced *α* occupancy (∼85%) compensated by an increased *β* content (∼10%). CHARMM22 and Amber ff03, which define the second generation, reveal more diverse conformational landscapes. In CHARMM22 simulations, most residues cluster tightly around *α*, *β*, and *ppII* propensities of ∼50%, ∼25%, and ∼25%, respectively, with glycine again being an outlier, favoring *ppII* (∼40%) at the expense of *β* structures (*<*10%). Amber ff03, on the other hand, shows a wider spread in *α* content: serine reaches values above 70%, while glutamic acid drops below 30%. The remaining amino acids fall between these limits, generally maintaining a near 1:1 ratio between *β* and *ppII*, except for serine and lysine, which show biases toward *α* and *ppII* conformations, respectively. The third generation of force fields, developed after 2017, reflects the concerted effort to improve the modeling of IDRs. Among these, CHARMM36m exhibits the lowest *α* content (10–25%) and a variable balance between *β* and *ppII*, aligning most closely with the GMM data for glycine. Amber ff19SB, which employs the 4-point OPC water model^1^, yields similar overall distributions, with most residues presenting 30–40% *α* and 50–65% *ppII* content, except glutamic acid, which shows higher *α* (over 50%) and lower *ppII* (around 40%). The force fields developed by the D.E. Shaw group (Amber ff99sb-disp and DESAmber) present highly consistent secondary-structure distributions across residues, with *ppII* around 50%, *α* between 20% and 30%, and *β* near 25%. Minor deviations occur for small residues such as glycine, serine, and leucine in ff99sb-disp, and glycine and serine in DESAmber, which display slightly enhanced *α* and reduced *ppII* propensities relative to the average.

### Wasserstein-based Multidimensional Scaling of (***ϕ***,***ψ***) Distributions of Guest Residues

In Supp. Figure 4, we report the two-dimensional scaling projection based on Wasserstein distances for each tripeptide, averaged over replicas. The overall picture is very similar to the two-dimensional scaling in Figure 2a. The oldest force field, ff94, is far from all other data sets, while second generation force fields (ff03 and charmm22) are close or corresponding to the PDB dataset, while the remaining force fields, belonging to the latest generation, are spread between the PDB-Exposed and the GMM data sets. Moreover, the three experimental data sets are often approximately aligned, indicating that by considering only the tripeptides of the PDB data set that are exposed to the solvent (*i.e.* the PDB-Exposed data set) leads to distributions that are more similar to those obtained with GMM. However, some notable differences appear. In the case of glycine as guest amino acid, even if the PDB and PDB-Exposed are clearly distinct, they have similar distances to the GMM data sets. This indicates that the distribution of (*ϕ, ψ*) obtained by considering only exposed tripeptides is not more similar to the GMM one. Nonetheless, also in the case of GGG, the distributions obtained from MD simulations (excluding ff94, which is always far from all other data sets) are still positioned on the imaginary line connecting PDB, PDB-Exposed, and GMM data sets. The distributions of PDB and PDB-Exposed data sets are more different (larger distance in the two-dimensional scaling) for smaller residues (GGG, GAG, and GSG) and more similar for amino acids with large or hydrophobic side chains (GFG, GLG, GEG, GKG).

## Supplementary Methods

### MD simulations

All Molecular Dynamics (MD) simulations were performed with Gromacs 2022^2^. The central amino acid set for the GXG tripeptides was chosen to be a good representative in terms of size, charge, and hydropathy (see Supp. Table 1). Simulations were performed using seven force fields from AMBER and CHARMM families developed in the last 30 years (see Supp. Table 2), namely AMBER ff94^3^, CHARMM22^4^, AMBER ff03^5^, CHARMM36m^6^, AMBER ff19SB^7^, amber99SB-disp^8^, and DES-Amber^9^. For each force field, the corresponding parameters for water and ions were used. For AMBER ff94, AMBER ff03 and CHARMM22 force fields, the parameters distributed with Gromacs 2022 were used. Parameters for CHARMM36m in Gromacs format were downloaded from http://mackerell.umaryland.edu/charmm_ff.shtml#gromacs (version charmm36-jul2020.ff). Parameters for AMBER ff99SB-disp and DES-Amber in Gromacs format were downloaded from https://github.com/paulrobustelli/Force-Fields. For simulations with the AMBER ff19SB force field, we generated a topology in AMBER format for each system using AmberTools LeAP utility and then converted them to Gromacs format using the amb2gro_top_gro.py python script included in AmberTools22^10^.

Initial configurations for the tripeptides were generated in extended configurations using the LEaP program included in the AmberTools22 package^10^. To avoid effects of the presence of terminal charges, we simulated all tripeptides with neutral termini: for simulations using parameters from the AMBER family, the termini of the tripeptides have been capped with NME and ACE groups, while for simulations of tripeptides using parameters from the CHARMM family, neutral NGLY and CGLY residues have been employed. Tripeptides have been solvated in rhombic dodecahedron boxes of ∼45.3 nm^3^ using a combination of gmx editconf and gmx solvate tools available in Gromacs 2022. The systems were then neutralized with 0.1 M NaCl using the gmx genion tool. Energy minimization with the steepest descent method was performed for up to 50000 steps to remove steric clashes between solvent and peptide molecules, followed by two rounds of equilibration of 100 ps each with a position restraint on the heavy atoms of the peptide with an harmonic constant set to 1000 kJ/mol/nm^2^, in the NVT and NPT ensembles, respectively. Temperature was set to 300 K and controlled using the vrescale algorithm^15^, while pressure was kept at 1 bar using the Parrinello-Rahman barostat^16^. Finally, a 1 *µ*s unrestrained MD simulation was run, saving peptide configurations every 10 ps for subsequent analysis. Bonds involving hydrogens have been constrained using LINCS^17^, allowing the use of 2 fs time steps for the integration of the equations of motion. Periodic boundary conditions were applied and long range electrostatic interactions were evaluated with the particle mesh Ewald algorithm ^18^ using a realspace cutoff of 1 nm. Van der Waals interactions were computed with a cutoff distance of 1 nm. For each peptide-force field combination, three independent replicas have been run.

**Supplementary Table 1.** : Properties of central amino acids used in this study, including van der Waals radii^11^ (in nm), net charge (q) of the side chain, and hydropathy (h) according to the Kyte-Doolittle scale^12^.

| Amino acid | vdW radius | q | h |
| --- | --- | --- | --- |
| Alanine (A) | 0.50 | 0 | 1.8 |
| Glutamic Acid (E) | 0.59 | -1 | -3.5 |
| Phenylalanine (F) | 0.64 | 0 | 2.8 |
| Glycine (G) | 0.45 | 0 | -0.4 |
| Lysine (K) | 0.64 | +1 | -3.9 |
| Leucine (L) | 0.62 | 0 | 3.8 |
| Serine (S) | 0.52 | 0 | -0.8 |

**Supplementary Table 2.**
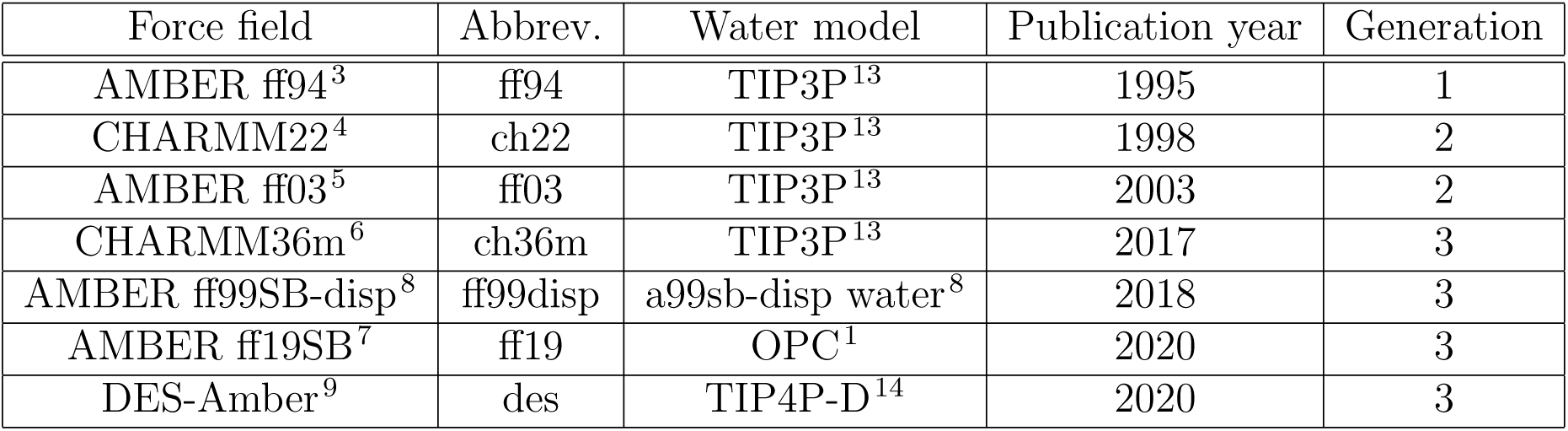
: Summary of the force fields employed in this study. Force fields are classified by generation, defined in the last column of the table.

| Force field | Abbrev. | Water model | Publication year | Generation |
| --- | --- | --- | --- | --- |
| AMBER ff94<sup>3</sup> | ff94 | TIP3P<sup>13</sup> | 1995 | 1 |
| CHARMM22<sup>4</sup> | ch22 | TIP3P<sup>13</sup> | 1998 | 2 |
| AMBER ff03<sup>5</sup> | ff03 | TIP3P<sup>13</sup> | 2003 | 2 |
| CHARMM36m<sup>6</sup> | ch36m | TIP3P<sup>13</sup> | 2017 | 3 |
| AMBER ff99SB-disp<sup>8</sup> | ff99disp | a99sb-disp water<sup>8</sup> | 2018 | 3 |
| AMBER ff19SB<sup>7</sup> | ff19 | OPC<sup>1</sup> | 2020 | 3 |
| DES-Amber<sup>9</sup> | des | TIP4P-D<sup>14</sup> | 2020 | 3 |

The Ramachandran dihedral angles, *ϕ* and *ψ*, of the central amino acids of tripeptides were evaluated using Plumed 2.8.0^19^. Convergence of simulations was evaluated by comparing the one-dimensional projections of the one-dimensional free energy profiles on the two Ramachandran dihedral angles of the central amino acid (see Supp. Fig. 2) between the independent replicas. Free energy profiles were obtained via Boltzmann inversion of the one-dimensional probability density functions *P* (*s*) as *F* (*s*) = −*k_B_T* ln *P* (*s*) + *C*, where *s* is either *ϕ* or *ψ*, *k_B_* is the Boltzmann constant, *T* is the temperature, and *C* is an arbitrary constant used to put the minimum of the free energy profile to 0.

### Definition of secondary structure regions

To assign the secondary structure of tripeptides, we employed the definition used by Best *et al.*^20^ and reported here in Supp. Table 3. Even if the choice of the definition of secondary structure regions is arbitrary, the qualitative balance of secondary structure populations is rather robust with respect to this choice. To this end, we recomputed the secondary structure populations using the definitions by Ting *et al.*^21^. We show the resulting triangular plot in Supp. Fig. 3. The average populations of all force fields, and particularly of all three generations is similar to what obtained with the definition by Best *et al.*, shown in Supp. Fig.1. Ternary plots with secondary structure populations have been plotted using the mpltern^22,23^ python library, version 1.0.4.

**Supplementary Table 3:** The three main secondary structure regions in the Ramachandran space, as defined by Best *et al.*.^24^

| Region | $\phi$ [deg] | $\psi$ [deg] |
| --- | --- | --- |
| $\alpha$ | $(-200, 0)$ | $(-120, 40)$ |
| $\beta$ | $(-90, 0) \cup (160, 180)$ | $(40, 240) \cup (110, 180)$ |
| $ppII$ | $[-200, 90)$ | $(40, 240]$ |

### Wasserstein-based Multidimensional Scaling

The Wasserstein distance represents the minimum transportation cost needed to reconfigure the mass of one probability distribution to recover the other. Formally, it can be computed as follows. If 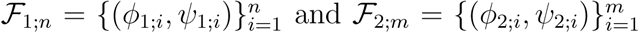 denote two samples drawn from a pair of dihedral angle distributions, their (2-)Wasserstein distance is given by:

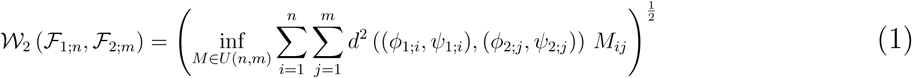

where *U* (*n, m*) is the set of real *n* × *m* matrices *M* = (*M_ij_*)*_ij_* such that 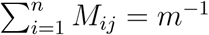 and 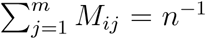, and *d* is the geodesic distance in the Ramachandran space. We refer to^25^ for an in-depth introduction to this distance in a more general context.

As the objective function and the constraints in Eq. (1) are linear in the variables of interest, the optimization problem is a linear program. Consequently, it can be solved with a large family of algorithmic tools from linear programming and combinatorial optimization. Here, we use the classical Network Simplex algorithm^26^ implemented in the R solver^27^.

To compare *N* samples of (*ϕ, ψ*) angles pairwise, we apply Eq. (1) to each pair, yielding an *N* × *N* matrix of Wasserstein distances. To visualize these results in a compact and informative manner, we employ Multidimensional Scaling (MDS)^28^. This algorithm projects the set of samples onto a two-dimensional Euclidean space, where the distances approximate the Wasserstein distances between the original samples as closely as possible. Here we used the implementation available in the R package stats^29^.

### PDB and PDB-Exposed datasets

As an experimental reference, we used a dataset composed of three-residue fragments (*i.e.*, tripeptides) extracted from coil regions of high-resolution structures in a non-redundant subset of the Protein Data Bank (PDB). This tripeptide dataset has been employed in previous studies to sample flexible and disordered protein regions ^30,31^ and to predict secondary-structure propensities within intrinsically disordered regions^32^. Further details about its construction can be found in these publications. It is worth noting that coil regions were defined using the DSSP classification^33^, which identifies secondary structure based on intra-backbone hydrogen-bonding patterns. As a consequence, tripeptides in this PDB-derived dataset may still adopt conformations characteristic of the *α*, *β*, or *ppII* classes, as these depend solely on their (*ϕ, ψ*) dihedral angles.

We also generated a more restrictive subset of the dataset by selecting only solvent-exposed tripeptides (PDB-Exposed). To do so, we applied a simple geometric filter: a tripeptide was discarded if more than five C*α* atoms from other residues were found within 8 Å of its middle residue’s C*α* atom. The number of data points for each GXG tripeptide in the data sets is reported in Supp. Table 4.

**Supplementary Table 4:** Number of data points for each GXG tripeptide in the PDB data set and in the PDB-exposed subset.

| GXG | PDB | PDB-Exposed |
| --- | --- | --- |
| GAG | 2366 | 446 |
| GEG | 1783 | 440 |
| GFG | 1363 | 209 |
| GGG | 2794 | 441 |
| GKG | 2000 | 557 |
| GLG | 2077 | 329 |
| GSG | 2315 | 410 |

### Gaussian mixture model

In the Gaussian Mixture Model (GMM)^34^, the probability distribution of (*ϕ, ψ*) angles is modeled as a mixture of 2-dimensional Gaussian distributions. Each mode corresponds to a different local secondary structure state, such as *ppII*, right- and left-handed *α*-helix, turns, parallel and antiparallel *β*, and various turns. In other words, the density function of (*ϕ, ψ*) is modeled by the function:

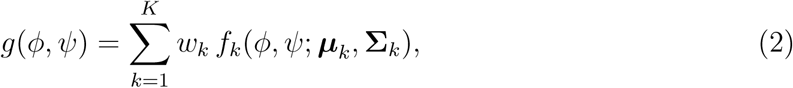

where, for each *k* = 1*, . . . , K*, *w_k_* is the weight of each component, *i.e.* the probability for an observation of being drawn from the *k*-th class, and *f_k_* is the density function of the *k*-th component, with parameters ***µ****_k_*, **Σ***_k_*. Each of the *f_k_* has the form:

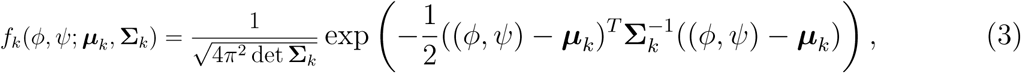

where ***µ****_k_* = (*µ_ϕ__k_, µ_ψ_*_;_*_k_*) is the mean vector of the *k*-th component and its covariance matrix **Σ***_k_* is diagonal with entries *σ_ϕ_*_;_*_k_* and *σ_ψ_*_;_*_k_*. The values of these parameters for every component and tripeptide are specified in Supp. Table 5.

The model parameters, given by the relative weights of the modes, together with their means and covariances, are adjusted to best fit the available J-coupling constants and amide I’ band profiles^34–37^. We present them in Supp. Table 5.

To compare the simulated distributions with the GMM, we sampled 100,000 (*ϕ, ψ*) pairs from the GMM of each GXG tripeptide, to match the size of the samples from each MD replica.

**Supplementary Table 5:** Parameters for Gaussian Mixture Model.

| Tripeptide | $\mu_{\phi;k}[\text{deg}]$ | $\mu_{\psi;k}[\text{deg}]$ | $\sigma_{\phi;k}[\text{deg}^2]$ | $\sigma_{\psi;k}[\text{deg}^2]$ | $w_k$ |
| --- | --- | --- | --- | --- | --- |
| GAG | -72 | 152 | 20 | 20 | 0.8 |
|  | -115 | 155 | 20 | 20 | 0.1 |
|  | -60 | -30 | 20 | 20 | 0.03 |
|  | -60 | 60 | 20 | 20 | 0.03 |
|  | 70 | -60 | 20 | 20 | 0.04 |
| GEG | -80 | 150 | 20 | 10 | 0.54 |
|  | -112 | 150 | 20 | 10 | 0.26 |
|  | -140 | 150 | 20 | 20 | 0.04 |
|  | -50 | -10 | 10 | 10 | 0.04 |
|  | 75 | 60 | 10 | 10 | 0.08 |
|  | 70 | 60 | 10 | 10 | 0.04 |
| GFG | -77 | 150 | 15 | 20 | 0.45 |
|  | -115 | 150 | 20 | 20 | 0.45 |
|  | -70 | -20 | 10 | 10 | 0.05 |
|  | 70 | 20 | 10 | 10 | 0.05 |
| GGG | -70 | 170 | 20 | 15 | 0.4 |
|  | 70 | -170 | 20 | 15 | 0.4 |
|  | -120 | 170 | 20 | 15 | 0.05 |
|  | 120 | -170 | 20 | 15 | 0.05 |
|  | -65 | -30 | 15 | 15 | 0.05 |
|  | 65 | 30 | 15 | 15 | 0.05 |
| GKG | -66 | 150 | 20 | 20 | 0.48 |
|  | -115 | 155 | 20 | 20 | 0.43 |
|  | -65 | -30 | 20 | 20 | 0.09 |
| GLG | -73 | 130 | 20 | 20 | 0.46 |
|  | -95 | 170 | 20 | 20 | 0.4 |
|  | -50 | -75 | 10 | 10 | 0.04 |
|  | 45 | -75 | 10 | 10 | 0.1 |
| GSG | -78 | 165 | 15 | 15 | 0.42 |
|  | -110 | 165 | 15 | 15 | 0.33 |
|  | -50 | 0 | 10 | 10 | 0.1 |
|  | 70 | 0 | 10 | 10 | 0.05 |
|  | 60 | 170 | 10 | 10 | 0.1 |

### Secondary structure proportions

**Supplementary Table 6:** Secondary structure propensities in the GMM, PDB and PDB-Exposed data sets.

| Data set | Amino acid | $\alpha$ | $\beta$ | $ppII$ | other |
| --- | --- | --- | --- | --- | --- |
| GMM | G | 0.100 | 0.099 | 0.801 | 0.000 |
|  | A | 0.031 | 0.099 | 0.830 | 0.041 |
|  | S | 0.099 | 0.332 | 0.419 | 0.150 |
|  | F | 0.052 | 0.452 | 0.447 | 0.049 |
|  | L | 0.040 | 0.350 | 0.510 | 0.100 |
|  | E | 0.040 | 0.310 | 0.533 | 0.117 |
|  | K | 0.091 | 0.429 | 0.481 | 0.000 |
| PDB | G | 0.654 | 0.177 | 0.157 | 0.012 |
|  | A | 0.474 | 0.178 | 0.239 | 0.109 |
|  | S | 0.555 | 0.184 | 0.172 | 0.089 |
|  | F | 0.318 | 0.337 | 0.194 | 0.151 |
|  | L | 0.531 | 0.185 | 0.177 | 0.107 |
|  | E | 0.354 | 0.266 | 0.263 | 0.117 |
|  | K | 0.364 | 0.264 | 0.247 | 0.125 |
| PDB-Exposed | G | 0.261 | 0.538 | 0.168 | 0.034 |
|  | A | 0.126 | 0.541 | 0.189 | 0.145 |
|  | S | 0.266 | 0.344 | 0.133 | 0.258 |
|  | F | 0.160 | 0.530 | 0.150 | 0.160 |
|  | L | 0.333 | 0.333 | 0.222 | 0.111 |
|  | E | 0.246 | 0.311 | 0.262 | 0.180 |
|  | K | 0.220 | 0.598 | 0.102 | 0.079 |

**Supplementary Table 7:** Secondary structure propensities in the MD simulations. Values are averaged over 3 independent replicas.

| Force field | Amino acid | $\alpha$ | $\beta$ | $ppII$ | Other |
| --- | --- | --- | --- | --- | --- |
| AMBER ff94 | G | 0.829 | 0.097 | 0.044 | 0.030 |
|  | A | 0.892 | 0.023 | 0.062 | 0.022 |
|  | S | 0.904 | 0.015 | 0.045 | 0.036 |
|  | F | 0.909 | 0.026 | 0.053 | 0.012 |
|  | L | 0.915 | 0.017 | 0.056 | 0.012 |
|  | E | 0.946 | 0.015 | 0.028 | 0.011 |
|  | K | 0.900 | 0.024 | 0.050 | 0.026 |
| CHARMM22 | G | 0.531 | 0.073 | 0.385 | 0.010 |
|  | A | 0.479 | 0.202 | 0.227 | 0.093 |
|  | S | 0.366 | 0.270 | 0.208 | 0.156 |
|  | F | 0.453 | 0.247 | 0.201 | 0.099 |
|  | L | 0.529 | 0.185 | 0.224 | 0.062 |
|  | E | 0.457 | 0.211 | 0.283 | 0.049 |
|  | K | 0.493 | 0.222 | 0.217 | 0.069 |
| AMBER ff03 | G | 0.313 | 0.334 | 0.349 | 0.004 |
|  | A | 0.422 | 0.216 | 0.283 | 0.078 |
|  | S | 0.681 | 0.142 | 0.119 | 0.058 |
|  | F | 0.378 | 0.285 | 0.284 | 0.053 |
|  | L | 0.461 | 0.220 | 0.278 | 0.041 |
|  | E | 0.258 | 0.393 | 0.294 | 0.056 |
|  | K | 0.596 | 0.131 | 0.237 | 0.035 |
| CHARMM36m | G | 0.119 | 0.102 | 0.778 | 0.001 |
|  | A | 0.218 | 0.221 | 0.503 | 0.059 |
|  | S | 0.159 | 0.326 | 0.403 | 0.113 |
|  | F | 0.178 | 0.277 | 0.500 | 0.045 |
|  | L | 0.219 | 0.236 | 0.489 | 0.056 |
|  | E | 0.174 | 0.183 | 0.596 | 0.047 |
|  | K | 0.218 | 0.240 | 0.484 | 0.057 |
| AMBER ff99sb-disp | G | 0.339 | 0.341 | 0.314 | 0.006 |
|  | A | 0.273 | 0.164 | 0.508 | 0.055 |
|  | S | 0.390 | 0.175 | 0.376 | 0.059 |
|  | F | 0.240 | 0.216 | 0.473 | 0.071 |
|  | L | 0.270 | 0.195 | 0.492 | 0.043 |
|  | E | 0.393 | 0.184 | 0.386 | 0.037 |
|  | K | 0.237 | 0.220 | 0.498 | 0.045 |
| AMBER ff19sb | G | 0.255 | 0.208 | 0.536 | 0.001 |
|  | A | 0.283 | 0.058 | 0.576 | 0.083 |
|  | S | 0.319 | 0.110 | 0.498 | 0.073 |
|  | F | 0.289 | 0.086 | 0.551 | 0.075 |
|  | L | 0.365 | 0.088 | 0.508 | 0.039 |
|  | E | 0.520 | 0.064 | 0.391 | 0.024 |
|  | K | 0.313 | 0.088 | 0.556 | 0.043 |
| DES-Amber | G | 0.340 | 0.362 | 0.296 | 0.002 |
|  | A | 0.233 | 0.208 | 0.504 | 0.055 |
|  | S | 0.292 | 0.320 | 0.326 | 0.061 |
|  | F | 0.173 | 0.305 | 0.486 | 0.037 |
|  | L | 0.267 | 0.267 | 0.431 | 0.035 |
|  | E | 0.280 | 0.263 | 0.427 | 0.030 |
|  | K | 0.178 | 0.302 | 0.493 | 0.026 |

**Supplementary Figure 1:**
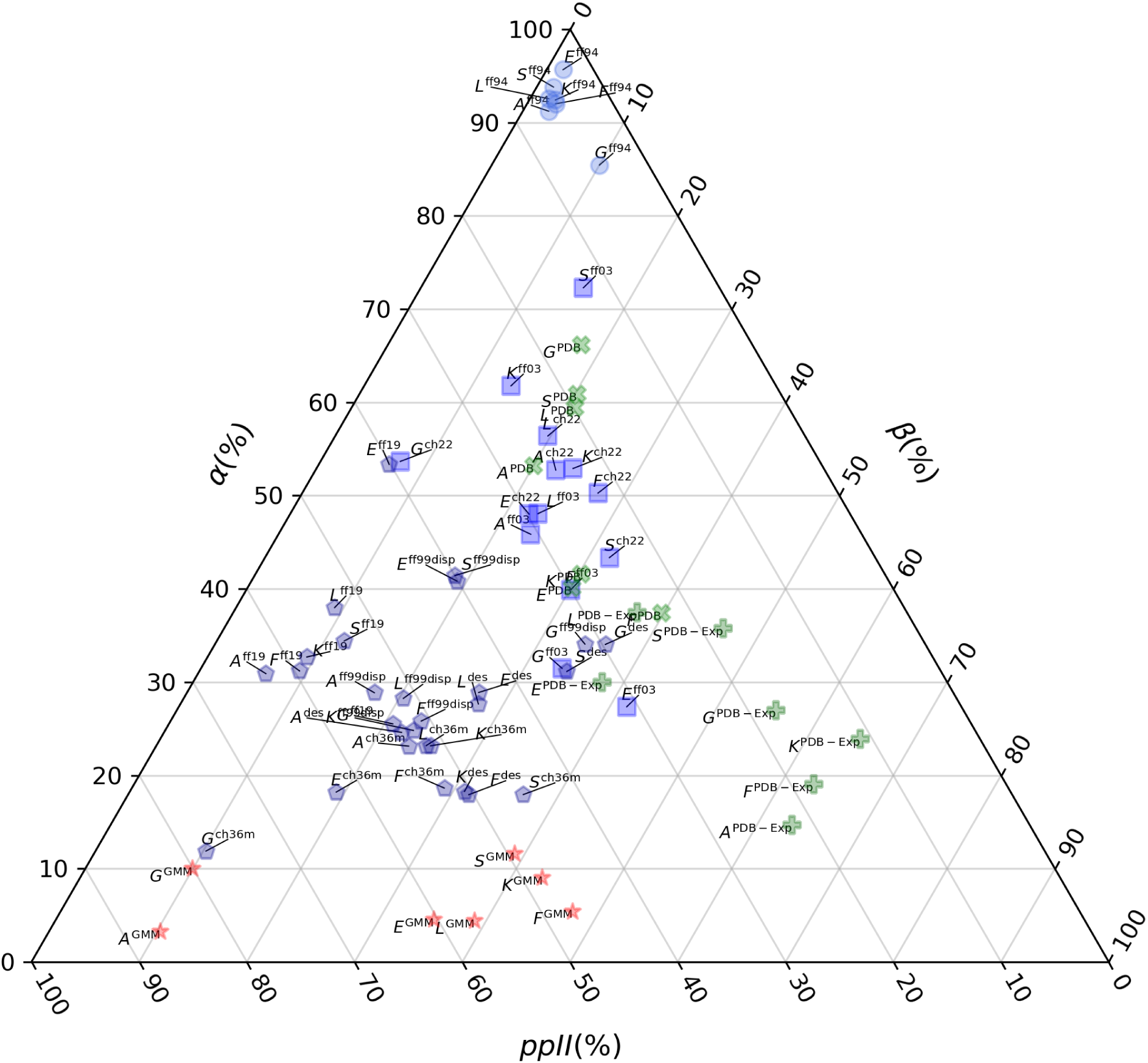
Triangular secondary structure populations for each tripeptide and force field using the definition by Best at al.^24^. Blue symbols indicate data from MD simulations, and the markers indicate the generation. Particularly, circles for the *first generation* (*i.e.* ff94), squares for the *second generation*, and pentagons for the *third generation*. Green symbols indicate data from PDB (× markers) and PDB-Exp (+ markers) data sets. Red stars indicate data from GMM data sets. Abbreviations for the force fields indicated as superscripts are reported in Supp. Table 2.

**Supplementary Figure 2:**
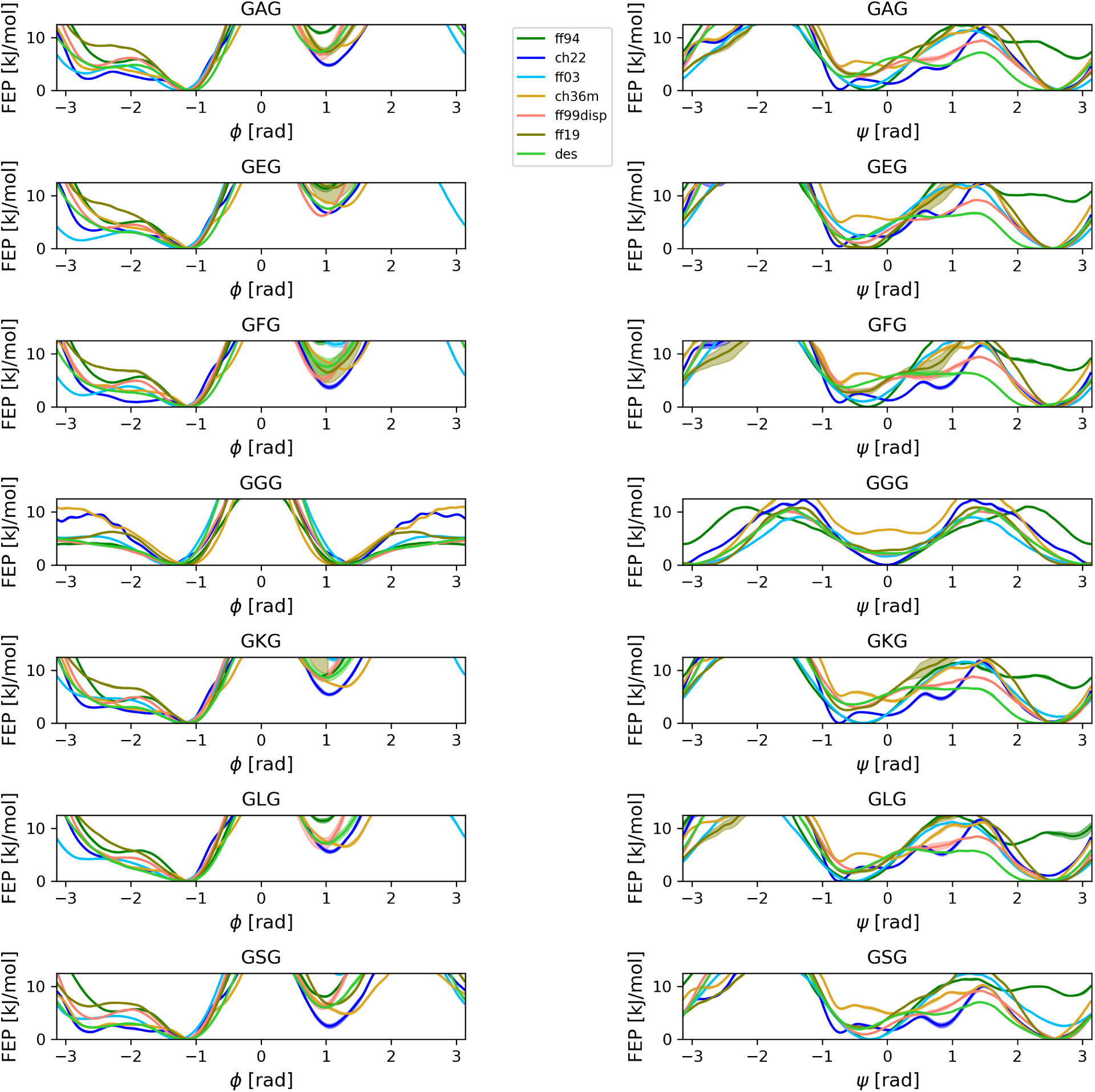
Free energy profiles as a function of Ramachandran angles (*ϕ*, *ψ*) from MD simulations for all central amino acids and force fields. Lines indicate averages over 3 independent replicas, shaded areas indicate standard deviation over the replicas. Abbreviations for the force fields are reported in Supp. Table 2.

**Supplementary Figure 3:**
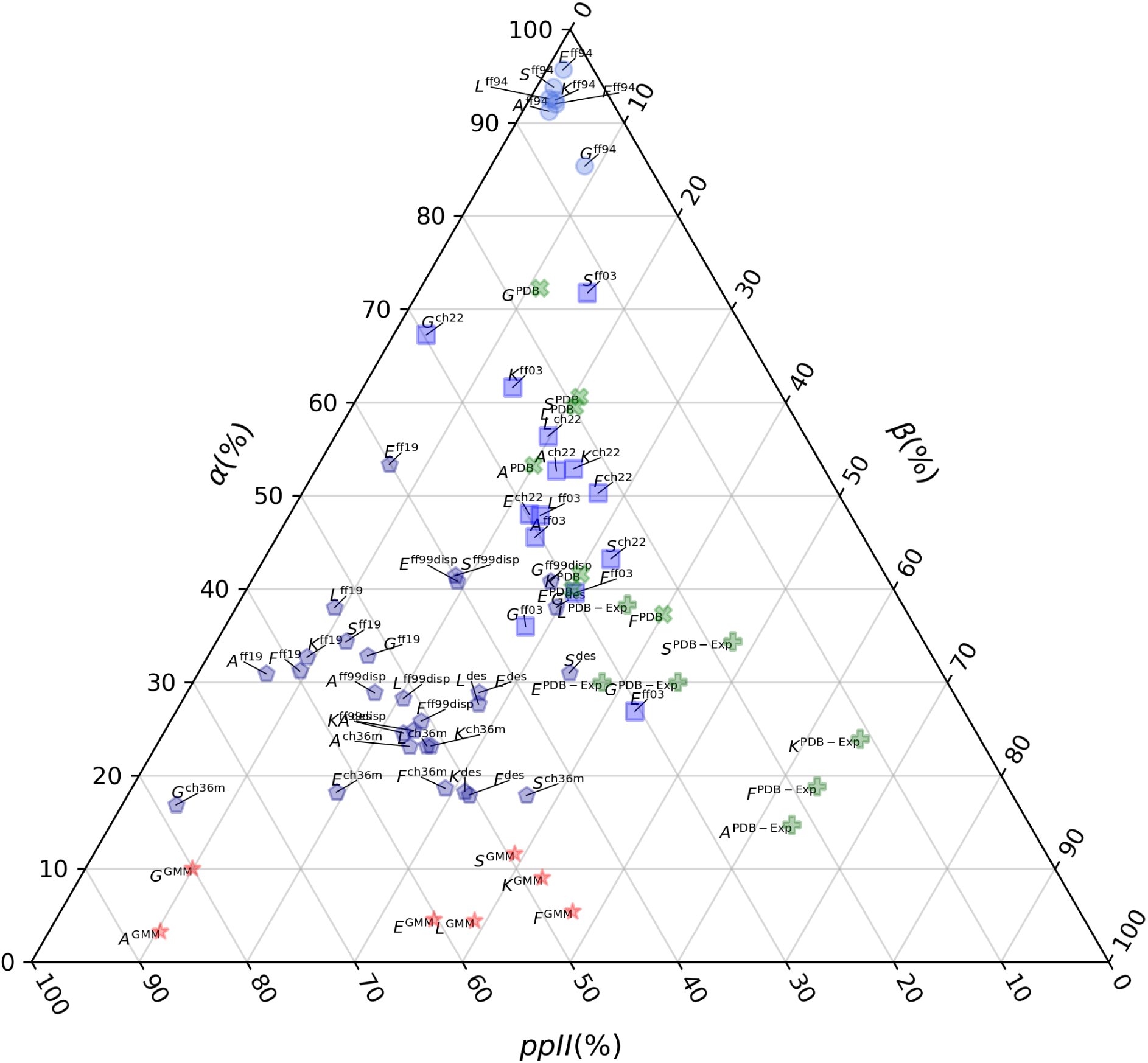
Triangular secondary structure populations for each tripeptide and force field using the definition by Ting *et al.*^21^. Blue symbols indicate data from MD simulations, and the marker indicate the generation. Particularly, circles for the *first generation* (*i.e.* ff94), squares for the *second generation*, and pentagons for the *third generation*. Green symbols indicate data from PDB (× markers) and PDB-Exp (+ markers) data sets. Red stars indicate data from GMM data sets. Abbreviations for the force fields indicated as superscripts are reported in Supp. Table 2.

**Supplementary Figure 4:**
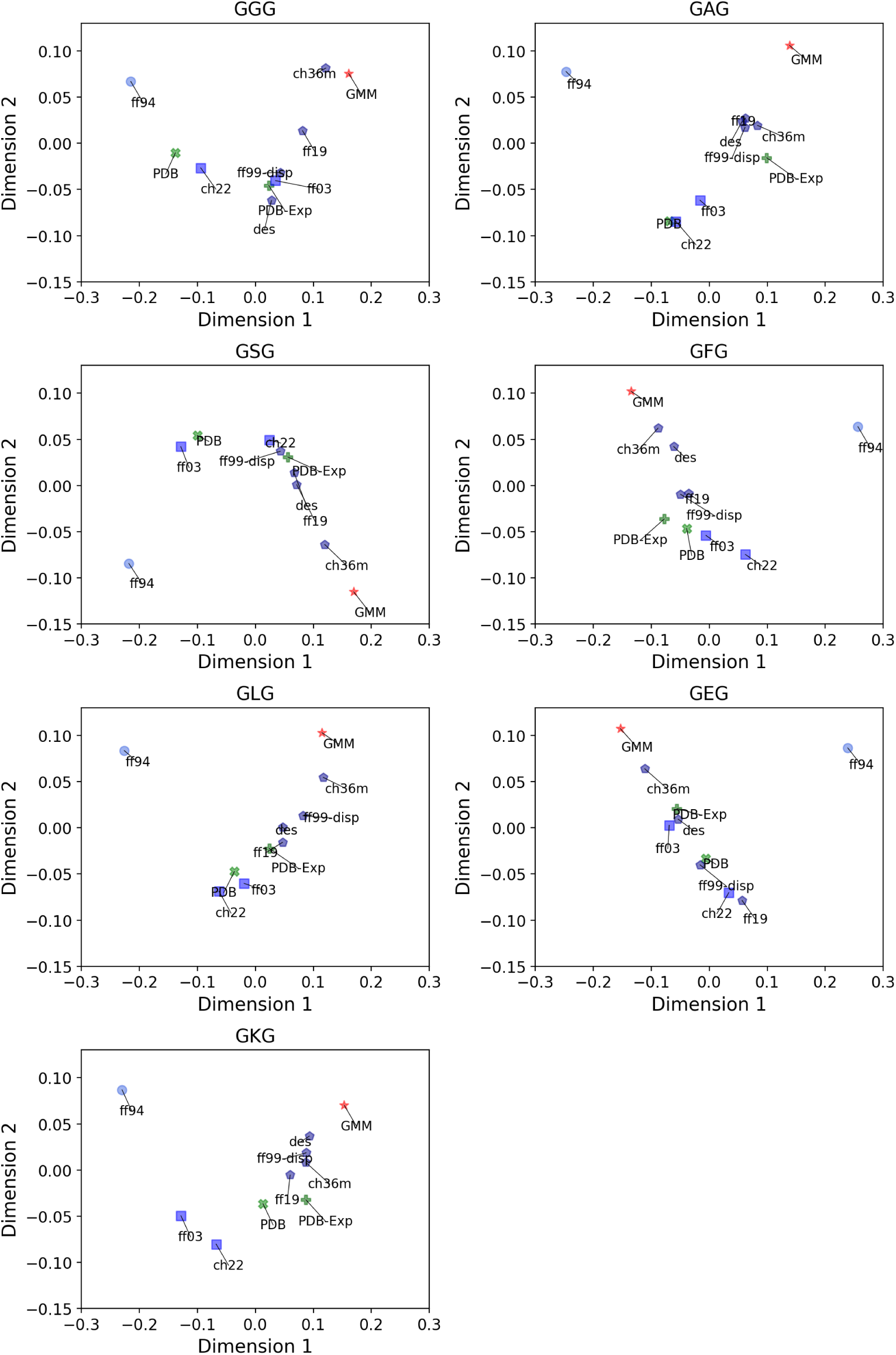
Wasserstein-based multidimensional scaling of (*ϕ*, *ψ*) distributions of each tripeptide. Blue symbols indicate data from MD simulations, and the marker indicate the generation. Particularly, circles for the *first generation* (*i.e.* ff94), squares for the *second generation*, and pentagons for the *third generation*. Green symbols indicate data from PDB (× markers) and PDB-Exp (+ markers) data sets. Red stars indicate data from GMM data sets. Abbreviations for the force fields are reported in Supp. Table 2.

## Notes

### Competing Interest Statement

The authors have declared no competing interest.

